# A Sense of Direction: Detecting Natural Selection Across Non-Stationary Genomes

**DOI:** 10.64898/2026.10.14.759254

**Authors:** Sergei L. Kosakovsky Pond

## Abstract

Codon substitution models detect natural selection by comparing non-synonymous (*dN*) and synonymous (*dS*) rates at each site. Standard methods assume stationary nucleotide composition across the phylogeny. In many genomes, however, GC-biased gene conversion (gBGC), codon usage bias, strand asymmetry, and host deaminases drive directional shifts in base composition along lineages. Under a stationary Muse–Gaut model with site-to-site synonymous rate variation (`FEL`), position-specific base frequencies and site-specific *dS* absorb most tree-wide drift. As a result, selection calls remain unchanged at 96%–99% of codons. At the remaining boundary sites, however, directional shifts bias *dS* in opposite directions depending on the ancestral codon. Rapid silent turnover at codons shifting toward the favored base inflates *dS*, producing apparent purifying selection and increasing across-site synonymous rate variance. Conversely, when amino acid replacements oppose the compositional tide, stationary models miss diversifying selection. Branch-specific models can accommodate shifting base frequencies, but diagonalizing a separate non-reversible codon rate matrix on every branch is too slow for large trees. Here we introduce Directional Fixed Effects Likelihood (`dFEL`). The method first tests whether base composition varies across taxa and runs standard `FEL` when frequencies are uniform. When composition shifts across lineages, `dFEL` roots the tree by maximum likelihood without an outgroup and models each branch as a smooth weighted average of three regime-specific transition probability matrices representing AT-enriching, neutral, and GC-enriching evolution. Sharing these matrices across the tree requires only three matrix diagonalizations per site. We validate `dFEL` across 1,800 simulated alignments and 87 unique empirical alignments (93 benchmark analyses) spanning up to 869 vertebrate species. It maintains calibrated false-positive rates, improves power, cuts whole-gene synonymous rate variance by up to 63%, and recovers established roots from coding sequences alone (https://github.com/veg/hyphy-3).

## Introduction

Codon substitution models detect natural selection in protein-coding genes by comparing the non-synonymous substitution rate (*β*_*s*_ or *dN*) with the synonymous rate (*α*_*s*_ or *dS*) at each codon site (Kosakovsky Pond and Frost, 2005; Murrell et al., 2012; Muse and Gaut, 1994). Because synonymous changes do not alter the amino acid, *α*_*s*_ serves as a local neutral baseline: sites with *β*_*s*_ *< α*_*s*_ evolve under purifying selection, whereas sites with *β*_*s*_ *> α*_*s*_ show diversifying positive selection. Standard implementations of Fixed Effects Likelihood (`FEL`) assume stationary base composition and time-reversibility across the phylogeny, so AT *→* GC and GC *→* AT fluxes are balanced at equilibrium on every branch and the tree likelihood is invariant to root placement (Felsenstein, 1981).

In many genomes, however, base composition shifts across lineages. In mammals and birds, GC-biased gene conversion (gBGC) favors G and C alleles during meiotic recombination repair, raising GC content on microchromosomes and in recombination hotspots (Bolívar et al., 2019; Duret and Galtier, 2009; Lartillot, 2013; Ratnakumar et al., 2010). In insects and bacteria, translational selection and asymmetric replication between leading and lagging DNA strands create strong third-position skews (Kreitman, 1983; McDonald and Kreitman, 1991). Similarly, viral host deaminases (APOBEC3 and ADAR) and vertebrate mitochondrial heavy-strand replication drive directional nucleotide shifts along specific clades.

How severely do these shifts affect site-level selection inference? Because standard `FEL` pairs a Muse– Gaut rate matrix (where substitution rates scale with the target nucleotide frequency rather than the full codon frequency) with a free site-specific synonymous rate *α*_*s*_ (Kosakovsky Pond and Frost, 2005; Muse and Gaut, 1994), it absorbs most tree-wide compositional drift without misclassifying conserved or neutral core sites. At boundary codons, however, unmodeled directional drift distorts 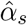 in two ways depending on the ancestral codon and the direction of amino acid change. In a clade shifting toward higher GC content, a functionally constrained site entering with A or T at the third position undergoes rapid synonymous AT *→* GC turnover. A stationary model interprets this silent turnover as a high synonymous rate 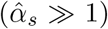, lowering 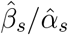 and producing apparent purifying selection. Conversely, at a site already carrying G or C at the third position, silent GC *→* AT turnover is suppressed 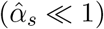, and when adaptive amino acid replacements move against the compositional tide, a stationary model loses power to detect diversifying selection. Splitting sites into rapid-turnover and silent-stasis classes also inflates apparent site-to-site synonymous rate variation (SRV) (Wisotsky et al., 2020).

Existing approaches to compositional heterogeneity face practical trade-offs. Branch-nonstationary models allow equilibrium frequencies to vary across branches (Boussau and Gouy, 2006; Galtier and Gouy, 1998), but diagonalizing a separate non-reversible 61 *×* 61 codon rate matrix on every branch at every site is too slow for large trees. Heuristic counting methods, such as the CpG-adjusted method of Suzuki et al. (2009) (SGK09), weight synonymous and non-synonymous opportunities by fixed mammalian mutation ratios and test parsimony counts with a Binomial distribution. However, discrete Binomial tests have low power on short alignments, and fixed mammalian CpG weights do not generalize to non-CpG biases.

Here we introduce Directional Fixed Effects Likelihood (`dFEL`), a fast extension of `FEL` in HyPhy 3 that accounts for lineage-specific GC and AT drift. `dFEL` first tests the alignment for compositional heterogeneity across taxa and runs standard `FEL` when base frequencies are uniform. When composition varies across lineages, `dFEL` roots the tree by maximum likelihood without an outgroup and models each branch as a smooth weighted average of three regime-specific transition probability matrices: AT-enriching, neutral, and GC-enriching. Sharing these three time-reversible generators across the tree requires only three matrix diagonalizations per site regardless of tree size. We validate `dFEL` on 1,800 simulated alignments and apply it to 87 unique empirical alignments (93 benchmark analyses) across classic benchmarks, avian macro- and microchromosomes, viral, mitochondrial, and bacterial strand-asymmetric loci, and deep vertebrate alignments (464–869 species) from the Vertebrate Genomes Project (VGP) and TOGA2.

## Model and Methods

Like standard `FEL` (Kosakovsky Pond and Frost, 2005), `dFEL` models substitutions among the 61 sense codons using a Muse–Gaut rate matrix (Muse and Gaut, 1994). At each codon site *s*, the model estimates a synonymous rate *α*_*s*_ and a non-synonymous rate *β*_*s*_, sharing global general time-reversible (GTR) nucleotide exchangeabilities *θ*_*uv*_ = *θ*_*vu*_ (*θ*_*AG*_ = 1) across the gene. Instead of one stationary frequency vector across the tree, `dFEL` builds three site-specific rate matrices 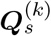: an AT-enriching regime (*k* = 0), a Neutral regime (*k* = 1), and a GC-enriching regime (*k* = 2). For codons *i* and *j* differing at a single codon position *m* ∈ {1, 2, 3} from base *u* to *v*, the instantaneous rate in regime *k* is given by Eq. (1).

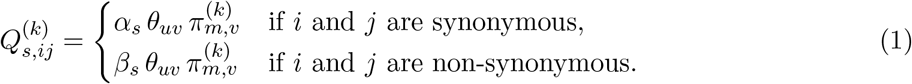

Multi-nucleotide changes have rate zero, and diagonal entries are 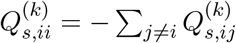.

The three regimes differ only in their target positional nucleotide frequencies 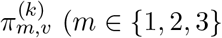, *v* ∈ {*A, C, G, T*}). Starting from alignment-wide empirical *F* 3 *×* 4 frequencies *π*_*m,v*_ (corrected for stop codons), `dFEL` tilts the frequencies in regime *k* by an exponential factor *ψ*^(*k*)^ (Eq. 2).

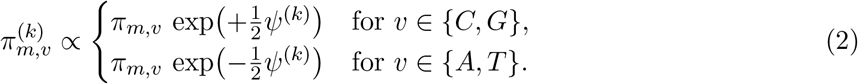

Rescaling the four bases at each position *m* to sum to 1 keeps 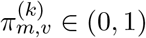 for any *ψ*^(*k*)^ and scales the AT *→* GC versus GC *→*AT rate ratio by exp(*ψ*^(*k*)^). Thus *ψ*^(*k*)^ is the log-odds shift in GC versus AT content relative to the alignment mean: *ψ*^(0)^ *<* 0 favors AT accumulation, *ψ*^(1)^ = 0 retains the baseline, and *ψ*^(2)^ *>* 0 favors GC accumulation. Equation (2) applies to all three codon positions (*m* = 1, 2, 3) because gBGC, strand skew, and deamination act on the DNA or RNA template regardless of reading frame; third positions absorb most of the shift at conserved sites (*β*_*s*_ *« α*_*s*_), whereas first and second positions also shift at unconstrained or positively selected sites.

Within each regime *k*, target codon frequencies ***π***^(*k*)^ are normalized products of positional frequencies 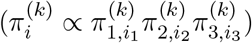. Root codon frequencies ***π***_*R*_ are fixed to the alignment baseline (***π***_*R*_ = ***π***^(1)^). Because each 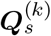 is time-reversible with respect to ***π***^(*k*)^, HyPhy 3 computes 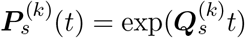 via symmetric eigendecomposition just like standard `FEL`.

Before fitting directional regimes, `dFEL` tabulates *N ×* 4 base counts across the *N* taxa for the full sequence and each codon position (CP1–CP3), evaluating each with a Pearson *χ*^2^ test (*df* = 3(*N −* 1)). If neither the full sequence nor CP3 is heterogeneous (*p ≥* 0.05), `dFEL` uses a single regime (*K* = 1) and returns the exact same estimates as standard `FEL`.

When heterogeneity is detected (*p <* 0.05), `dFEL` activates all three regimes (*K* = 3) in a two-stage plug-in procedure. In Phase 1, `dFEL` fits a nucleotide GTR model for initial branch lengths and *θ*_*uv*_, assigns branch weights *w*_*b,k*_ and regime tilts *ψ*^(*k*)^, scans candidate roots, and fits a global codon rate scalar and *ω*. Because branches with different compositional trends use different transition matrices while the root uses ***π***_*R*_, the tree likelihood depends on the root node *R*. To root the tree without an outgroup (`--root auto`), `dFEL` places a trial root at the midpoint (0.5*t*_*b*_) of each of the 2*N −* 3 branches, selects the branch maximizing ln *L*(*b*), and summarizes support across equal-parameter candidate roots via relative likelihood weights, *w*_root_(*b*) *∝* exp(ln *L*(*b*) *−* ln *L*_max_). When an outgroup root is known, `--root keep` skips this scan.

Rather than assigning branches to discrete regimes with a hard GC_3_ cutoff, `dFEL` models each branch *b* (leading to node *u*) as a smooth mixture of the three regimes (Fig. 1A). For each node *u*, it computes the third-position GC deviation 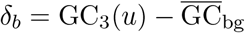 (using the descendant-tip mean GC_3_ for internal nodes). Branches near zero weight the Neutral regime (*w*_*b*,1_ *≈* 1), whereas AT-shifted (*δ*_*b*_ *<* 0) and GC-shifted (*δ*_*b*_ *>* 0) branches shift toward AT-drift (*w*_*b*,0_) or GC-drift (*w*_*b*,2_) via Softmax weights (Eq. 3).

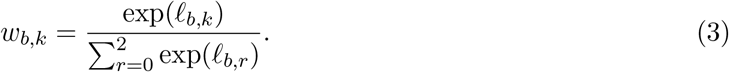

**Figure 1.**
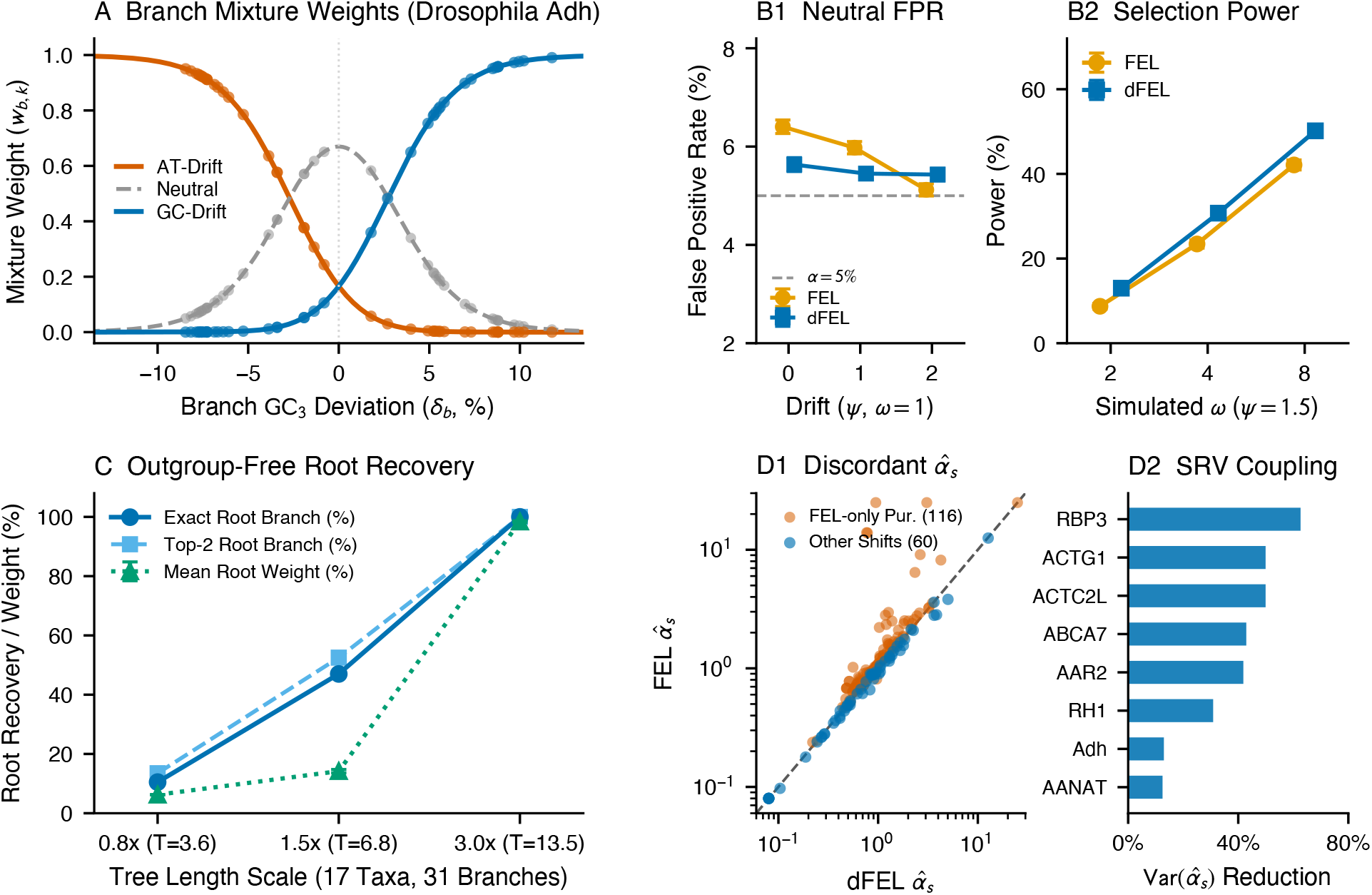
Model weights, simulation validation, and coupling between directional drift and synonymous rate variation (SRV). A, Branch mixture weights *w*_*b,k*_(*δ*_*b*_) across the 43 branches of *Drosophila Adh*, showing the transition from the maximum-likelihood root on *D. picticornis* (pic) to the GC-rich *melanogaster* clade. B1, Neutral false-positive rate (*ω* = 1.0) across *ψ ∈ {*0, 1, 2} and B2, positive selection sensitivity (*ψ* = 1.5) across *ω ∈ {*2, 4, 8} for FEL (amber) and dFEL (blue) across 200 replicates per condition (markers: mean *±*1 SEM). C, Outgroup-free root recovery accuracy (exact root branch, top-2 branch, and replicate relative likelihood root weights) across tree divergence scales (0.8*×*, 1.5*×*, 3.0*×*; 200 replicates per scale, 17 taxa, 31 branches). D1, Site-level 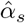 estimates across the 176 discordant sites in the avian and asymmetry benchmark suites (FEL-only purifying calls vs. other shifts) and D2, percent reduction in whole-gene across-site Var 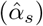 under dFEL.

Here *l*_*b*,0_ = *κ*(*−δ*_*b*_ *− τ*), *l*_*b*,1_ = 0, and *l*_*b*,2_ = *κ*(*δ*_*b*_ *− τ*), where *τ* = max(0.02, 0.4*σ*) and *κ* = 3.5*/σ* scale with the across-branch standard deviation *σ* of *δ*_*b*_ (*σ ≥* 0.015). From the weighted mean GC_3_ in each regime 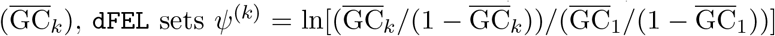, bounded to [*−*3.0, *−*0.05] (*k* = 0) and [0.05, 3.0] (*k* = 2).

The transition probability matrix along branch *b* of length *t*_*b*_ at site *s* is a convex combination of the three regime transition matrices (Eq. 4).

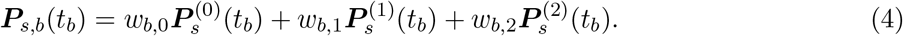

Combining matrices at the transition-probability level allows all branches to share the same three eigendecompositions 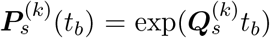, requiring only three matrix diagonalizations per site. In Phase 2, with Phase 1 parameters fixed, `dFEL` estimates *α*_*s*_ and *β*_*s*_ at each site *s* by maximum likelihood under *H*_1_ : *α*_*s*_ = *β*_*s*_ and *H*_0_ : *α*_*s*_ ≠ *β*_*s*_, testing significance via an asymptotic 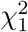 likelihood ratio test (LRT_*s*_ = 2(ln *L*_1_ *−* ln *L*_0_)), and reconstructs ancestral codons (Pupko et al., 2000) to record directed substitutions.

## Results

We first evaluated `dFEL` against stationary `FEL` across three simulation suites totaling 1,800 alignments (200 replicates per condition, 150 codons, 17 taxa [16 ingroup plus 1 outgroup], 31 branches, total tree length *T* = 4.5 substitutions per site; Fig. 1B1–C). Under strictly neutral evolution (*ω* = 1.0; Fig. 1B1), directional drift acts across all three codon positions and scales synonymous and non-synonymous rates together (*α*_*s*_ = *β*_*s*_ = 1). Because all three codon positions drift together when amino acids are unconstrained, stationary `FEL` does not accumulate apparent purifying calls under *ω* = 1.0. In empirical proteins, by contrast, purifying constraint holds first and second positions fixed while third positions turn over rapidly. Both methods stay near the nominal *α* = 0.05 level across all drift strengths (`FEL`: 5.1%–6.4%; `dFEL`: 5.4%–5.6% for *ψ ∈ {*0, 1, 2}). Directional regimes therefore do not inflate Type I error.

Lineage-specific compositional drift reduces the power of stationary models when adaptive amino acid changes occur against a shifting background. When diversifying selection acts on 6% of codons (*ω ∈ {*2, 4, 8}) under directional drift (*ψ* = 1.5; Fig. 1B2), `dFEL` increases sensitivity by 4.3 to 8.1 percentage points over stationary `FEL` (13.0% vs. 8.7% at *ω* = 2; 30.7% vs. 23.4% at *ω* = 4; 50.2% vs. 42.1% at *ω* = 8). Root identifiability across the 31 branches of the unrooted 17-taxon tree also rises with sequence divergence (Fig. 1C). Exact root recovery climbs from 10.5% on shallow trees (0.8*×*) to 47.0% at intermediate depth (1.5*×*) and reaches 100% on deep trees (3.0*×*, mean root weight 98.4%).

We next compared `FEL` and `dFEL` with the CpG-adjusted counting method of Suzuki et al. (2009) (SGK09; Supplementary Table S3). SGK09 tests whether positive selection calls in primate Protamine 1 (*PRM1*; 24 taxa, 48 codons) reflect CpG hypermutation in CGN arginine codons. It weights opportunities by fixed mammalian CpG mutation ratios and applies a parsimony Binomial test.

Because this weighting raises the expected non-synonymous fraction at CGN codons to 0.827, reaching *p ≤* 0.10 requires at least 13 non-synonymous changes and zero synonymous changes at a single site. On short alignments, this discrete cutoff detects zero selected sites in *PRM1* and misses codon 197 in *Drosophila Adh* (*p* = 0.569). By contrast, both `FEL` and `dFEL` (which are identical homogeneous *PRM1, K* = 1) detect diversifying selection at codons 20 and 45 (*p ≤* 0.05; plus codon 7 at *p ≤* 0.10) and recover codon 197 in heterogeneous *Adh* (*K* = 3, *p* = 0.015). Conversely, under non-CpG gBGC in avian *ABHD14A* (*K* = 1) and *AAR2* (*K* = 3), fixed mammalian CpG weights mismatch the substitution bias and yield three and eight excess positive selection calls (*p ≤* 0.05) unsupported by `FEL` or `dFEL`.

Comparing site-level 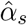 estimates between `FEL` and `dFEL` on empirical data shows how directional base-composition shifts inflate apparent SRV under stationary models (Fig. 1D1–D2). When a clade shifts toward high GC content at a constrained site, codons entering with A or T at silent positions undergo rapid AT *→* GC turnover 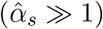, whereas codons already carrying G or C remain static 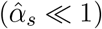. Across the 176 discordant sites in our avian and asymmetry suites, stationary `FEL` estimates a higher synonymous rate 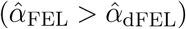 at 95.7% (111/116) of sites called purifying by `FEL` only (Fig. 1D1). Accounting for directional regimes in `dFEL` cuts Var 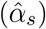 by 88.5% across discordant sites (12.53 *→* 1.44) and by 12.5%–62.6% across whole genes with strong GC_3_ divergence, including *RBP3* (*−*62.6%), *ACTG1* and *ACTC2L* (*−*49.9%), *ABCA7* (*−*42.9%), *AAR2* (*−*41.8%), and *RH1* (*−*30.9%; Fig. 1D2; Supplementary Table S4).

### Empirical Applications

We applied `dFEL` across 87 unique empirical alignments (93 benchmark analyses; six reference loci appear in both the classic and asymmetry suites) spanning four settings (Fig. 2, Table 1, and Supplementary Tables S1–S4). Across all 32 compositionally homogeneous (*K* = 1) alignments in the four suites, `FEL` and `dFEL` produced 100% identical site calls (0 discordant sites).

**Table 1.** Summary of dFEL results across the empirical benchmark suites (87 unique alignments / 93 benchmark analyses; Supplementary Tables S1–S4). Site differences are reported at *p ≤* 0.05 (or *p ≤* 0.10 where noted).

| Dataset / Cohort | Taxa | Codons | CP3 $\chi^2$ ($p$) | CP3 GC | Auto-Root | $w(b)$ | $K$ | Time | Site Call & SRV Differences |
| --- | --- | --- | --- | --- | --- | --- | --- | --- | --- |
| *Drosophila Adh* | 23 | 254 | 288 ($10^{-29}$) | 59–85% | pic | 4.6% | 3 | 3.6 s | Cuts $\text{Var}(\hat{\alpha}_s)$ by 13.0%; detects Div site 197 (4 shifts at $p \leq 0.10$). |
| Avian *AAR2* | 26 | 336 | 342 ($10^{-37}$) | 54–91% | taeGut2 | 18.2% | 3 | 3.8 s | 13 Pur in FEL only (120 → 107); cuts $\text{Var}(\hat{\alpha}_s)$ by 41.8%. |
| Avian gBGC (20 loci) | 22–44 | 10,091 | 10 $K = 1$, 10 $K = 3$ | 26–97% | Outgroup-free | — | 1–3 | 3–58 s | 0 diff on 10 $K = 1$ loci; 95 shifts on 10 $K = 3$ loci (+4 Div on micro). |
| Primate *PRM1* | 24 | 48 | 7.7 (1.00) | 44–65% | Homog. | — | 1 | 0.8 s | dFEL = FEL: both detect 2 Div / 1 Pur (3/2 at $p \leq 0.10$) vs. SGK09 (0/0). |
| Asymmetry (33 loci) | 10–180 | 14,194 | 9 $K = 1$, 24 $K = 3$ | 4–87% | Outgroup-free | — | 1–3 | 1–24 s | 0 diff on 9 $K = 1$ loci; 81 shifts on 24 $K = 3$ loci (48 FEL-only Pur). |
| Vertebrates (24 VGP) | 464–869 | 5,923 | 14 $K = 1$, 10 $K = 3$ | 18–92% | eleEdw1 | — | 1–3 | 2–5 m | 0 diff on 14 $K = 1$ loci; 26 shifts on 10 $K = 3$ loci ($p \leq 0.10$). |
| SARS-CoV-2 Spike | 180 | 1,284 | 4.4 (1.00) | 26–27% | Homog. | — | 1 | 22.1 s | Homogeneous ($K = 1$): 144 Pur / 21 Div, 0 discordant sites. |

**Figure 2.**
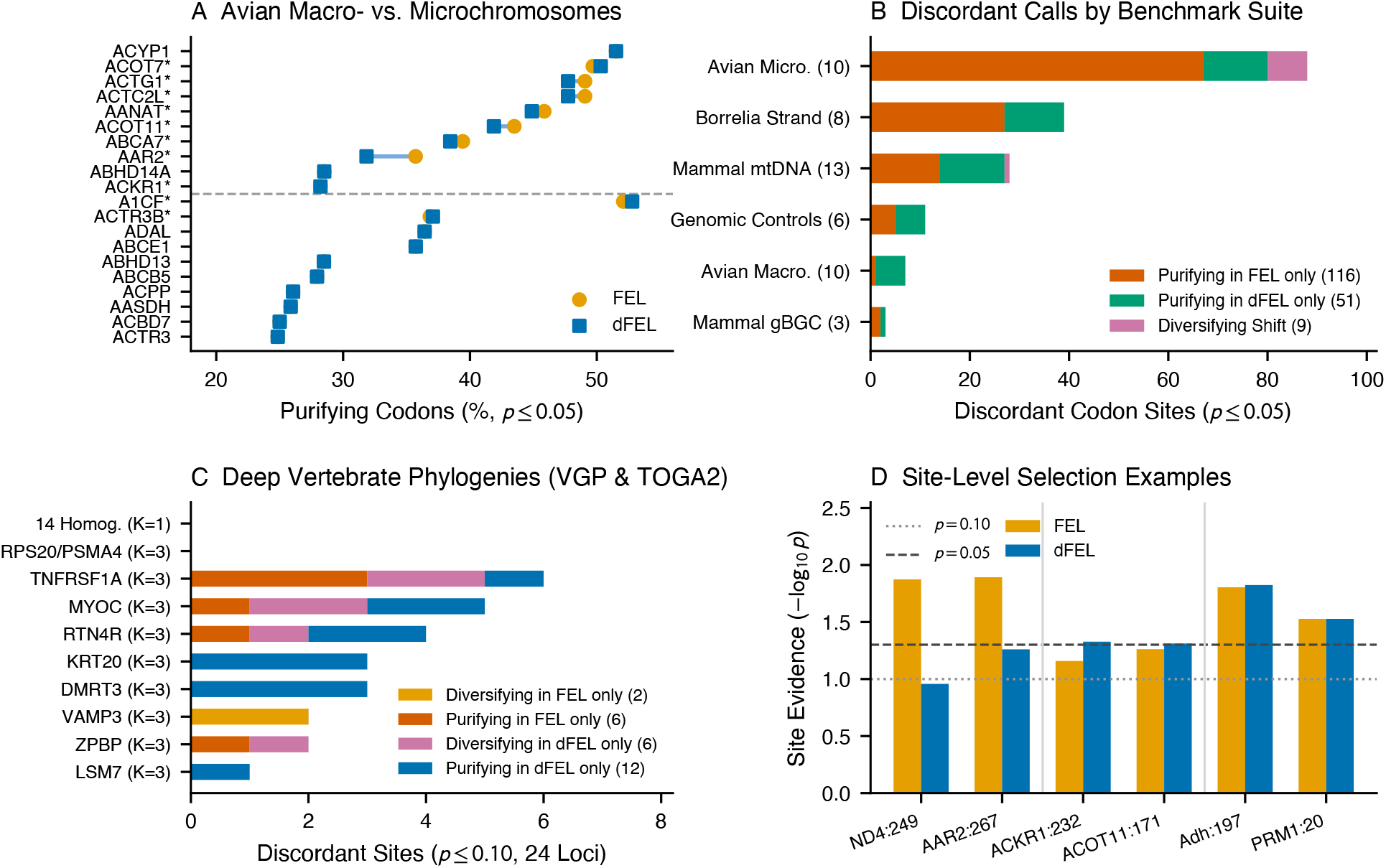
Site-level selection calls under FEL and dFEL across 87 unique empirical alignments (93 benchmark analyses). A, Percentage of codons classified as purifying (*p ≤*0.05) by FEL (amber circles) and dFEL (blue squares) across 10 microchromosomal (top) and 10 macrochromosomal (bottom) avian genes; asterisks denote heterogeneous (*K* = 3) loci. B, Breakdown of 176 discordant codon sites (*p≤* 0.05) across the avian micro- and macrochromosome suites and the 33-locus asymmetry benchmark. C, Discordant site counts (*p≤*0.10) across 24 deep vertebrate alignments (464–869 species, 5,923 codons), contrasting 14 homogeneous (*K* = 1) genes with 10 heterogeneous (*K* = 3) genes. D, Selection evidence (*−* log_10_ *p*) at six representative codons showing tempered purifying calls (*ND4* :249, *AAR2* :267), unmasked diversifying calls (*ACKR1* :232, *ACOT11* :171), and concordant diversifying calls missed by SGK09 (*Adh*:197, *PRM1* :20).

In *Drosophila Adh* (23 species, 254 codons; Fig. 1A), third-position GC rises from 58.7% in *D. picticornis* to 85.4% in *D. melanogaster*. Without an outgroup, `dFEL` rooted the tree on *D. picticornis*, capturing the shift from an AT-rich ancestor to the GC-rich *Sophophora* clade, cutting whole-gene Var 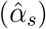 by 13.0%, and recovering diversifying selection at codon 197 (*p* = 0.015; Fig. 2D).

A broader test of gBGC spans 20 avian genes from macrochromosomes (10 genes, 5,031 codons) and microchromosomes (10 genes, 5,060 codons; Fig. 2A–B and Supplementary Table S1), where elevated recombination on microchromosomes drives strong AT *→* GC conversion (Bolívar et al., 2019). Ten avian genes (8 macrochromosomal and 2 microchromosomal, including uniformly high-GC *ABHD14A*, GC_3_ = 73%–91%) are stationary across lineages (*K* = 1) and show zero discordance. The 10 heterogeneous genes (*K* = 3: 2 macrochromosomal, 8 microchromosomal) contain 95 discordant sites (*p ≤* 0.05). On the 8 heterogeneous microchromosomes, rapid AT *→* GC silent turnover inflates stationary 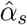 at conserved AT-ending codons. Modeling directional drift in `dFEL` tempers 67 `FEL`-only purifying calls while unmasking 13 `dFEL`-only purifying calls (net reduction from 1, 954 [41.4%] to 1, 900 [40.3%] purifying codons, e.g., *AAR2* codon 267: *p*_FEL_ = 0.013 *→ p*_dFEL_ = 0.055). It also increases diversifying calls on microchromosomes from 15 to 19 (+4 sites: *ABCA7* 9 *→* 11, *ACKR1* codon 232 [*p*_FEL_ = 0.069 *→ p*_dFEL_ = 0.047], and *ACOT11* codon 171 [*p*_FEL_ = 0.055 *→ p*_dFEL_ = 0.049]; Fig. 2D).

We next examined 33 alignments with mutational skews across mammalian mitochondria (13 genes), *Borrelia* replication strands (8 genes), mammalian gBGC hotspots (3 genes), viral APOBEC3 and ADAR targets (3 genes), and six controls (14,194 codons; Fig. 2B and Supplementary Table S2).

Nine homogeneous alignments (*K* = 1, including viral deamination and SARS-CoV-2 Spike) showed zero discordant sites. Across the 24 heterogeneous alignments (*K* = 3), `FEL` and `dFEL` differed at 81 sites (*p ≤* 0.05): 39 in *Borrelia*, 28 in mammalian mtDNA, 11 in heterogeneous controls, and 3 in mammalian gBGC loci. Stationary `FEL` called 48 of these sites purifying that became neutral under `dFEL` once lineage-specific silent turnover was modeled (e.g., mammalian *ND4* codon 249, where 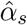 dropped from 13.89 to 0.78 and *p* shifted from 0.013 to 0.110; Fig. 2D), whereas `dFEL` unmasked 32 purifying sites where silent stasis had depressed 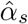 under `FEL`. On unrooted inputs, the root scan also recovered established basal lineages from coding sequences alone, rooting mammalian mtDNA on the platypus (*Ornithorhynchus anatinus*) and bacterial *rpoB* on *Candidatus Borrelia tachyglossi*.

Finally, across 24 deep vertebrate alignments from VGP and TOGA2 (464–869 species, up to 1,521 branches; Fig. 2C and Supplementary Table S4), we evaluated calls at *p ≤* 0.10 (the standard HyPhy Vision threshold for exploratory comparative scans). `FEL` and `dFEL` agreed at 99.56% of all 5,923 codons, with 100% concordance across all 14 homogeneous genes (*K* = 1, 2,844 codons). All 26 discordant sites occurred in the 10 heterogeneous (*K* = 3) genes: 6 purifying and 2 diversifying in `FEL` only, versus 12 purifying and 6 diversifying in `dFEL` only. At *VAMP3* codon 94, `FEL` inferred diversifying selection (*p* = 0.089) across 15 recurrent ATT/ATC *→* GTT/GTC (I *→* V) changes in high-GC mammals; `dFEL` attributed these GC-increasing changes to lineage drift and classified the site as neutral (*p* = 0.277), matching episodic `MEME` (*p* = 0.372). Conversely, `dFEL` identified diversifying selection at *MYOC* codon 15 (*p*_FEL_ = 0.102 *→ p*_dFEL_ = 0.085), corroborated by `MEME` (*p* = 0.048).

Scanning all candidate roots (`--root auto`) on six deep trees produced the exact same site calls as fixing the species-tree root (`--root keep`) and placed the root on Afrotheria (*Elephantulus edwardii*) in *ADORA3* and *WDR83OS*.

## Discussion

A central finding of our benchmarks is that standard `FEL` and directional `dFEL` agree on the selective classification of 96%–99% of codon sites across compositionally heterogeneous genes. Why is the standard reversible approach so resilient to directional base-composition drift? Unlike Goldman–Yang codon models (Goldman and Yang, 1994), where rates scale with full target codon frequencies, `FEL` uses the Muse–Gaut rate matrix (Muse and Gaut, 1994) with position-specific nucleotide frequencies (F3 *×* 4) and a free site-specific synonymous rate *α*_*s*_ (Kosakovsky Pond and Frost, 2005). Because the directional tilt exp 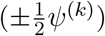 scales synonymous and non-synonymous changes toward the same target base by the same factor, and because *α*_*s*_ absorbs tree-wide differences in silent substitution opportunity at each site, stationary `FEL` remains well calibrated at most sites. Modeling directional drift is needed in three specific settings. First, it recalibrates the 1%–4% of boundary codons where rapid AT *→* GC silent turnover at conserved sites inflates 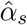 and whole-gene Var 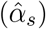 by up to 62.6% (Wisotsky et al., 2020), or where amino acid replacements oppose the compositional tide. Second, it infers outgroup-free maximum-likelihood roots and directed substitution histories directly from coding sequences. Third, it provides a fast three-eigenvalue mixture framework for episodic branch-site models such as `MEME` (Murrell et al., 2012) and `BUSTED` (Murrell et al., 2015), where a single site-wide *α*_*s*_ cannot absorb branch-specific bursts of compositional drift.

`dFEL` captures these directional effects without diagonalizing a separate rate matrix on every branch. Its *χ*^2^ homogeneity gate guarantees zero divergence from standard `FEL` when base composition is stationary (*K* = 1), whereas heterogeneous alignments activate the three-regime mixture (*K* = 3) using only three matrix diagonalizations per site (Kosakovsky Pond et al., 2020). Two approximations point to natural extensions. First, `dFEL` tilts nucleotide frequencies along a single AT *↔*GC axis (*ψ*^(*k*)^); in replicons with strong G*/*T strand skews (such as *Borrelia* chromosomes and vertebrate mitochondria), *ψ*^(*k*)^ captures the net GC_3_ shift, and a second tilt axis (*ψ*_GT_) would accommodate within-class skews. Second, fixing root codon frequencies ***π***_*R*_ to the alignment average avoids extra free parameters during root scanning across all 2*N −* 3 branches, though basal branches absorb the offset when taxon sampling is heavily skewed (as on the *D. picticornis* branch in *Adh*). Because site-level selection calls are invariant to root placement on deep vertebrate trees, users analyzing rooted phylogenies can fix the input root (`--root keep`) to cut runtime by three-to five-fold, reserving `--root auto` for unrooted trees.

## Software and Data Availability

`dFEL` is implemented in C++20 in HyPhy 3. It runs from the command line (`hyphy3 dfel` or `hyphy3-dfel`) or from Python (`import hyphy3 as hp`), and writes JSON output compatible with HyPhy Vision (https://vision.hyphy.org). All benchmark alignments, source code, and tests are available under the MIT license at https://github.com/veg/hyphy-3.

