## Supplementary Text and Tables for "A Sense of Direction: Detecting Natural Selection Across Non-Stationary Genomes"

### Supplementary Note: Empirical Benchmarking Across Mechanistic Asymmetry

Stationary codon substitution models assume that equilibrium base frequencies stay constant across all branches of a phylogenetic tree. Under this assumption, directional mutation biases—such as deamination, strand-asymmetric replication error, or GC-biased gene conversion—are confounded with natural selection. When nucleotide mutation rates are asymmetric, stationary models absorb directional nucleotide changes into an inflated synonymous rate estimate $\hat{\alpha}_s$ at sites entering a drifting clade in an unfavored nucleotide state, while depressing $\hat{\alpha}_s$ at sites already fixed in the favored nucleotide state. Stationary `FEL` then inflates across-site synonymous rate variance $\mathrm{Var}(\hat{\alpha}_s)$, computes a distorted ratio $\hat{\omega}_s = \hat{\beta}_s/\hat{\alpha}_s$, and misclassifies neutral sites as purifying.

To test this prediction on empirical data, we compared stationary `FEL` and directional `dFEL` across 33 alignments with known compositional asymmetry (Supplementary Table S2). We also evaluated 24 deep vertebrate alignments (464–869 species) from the Vertebrate Genomes Project (VGP) and TOGA2 (Supplementary Table S4):

1. **Vertebrate Mitochondrial DNA (13 protein-coding genes):** Asymmetric replication of the mammalian mitochondrial genome leaves the heavy strand single-stranded during replication. Spontaneous cytidine and adenine deamination creates A/C-rich heavy-strand genes and an inverted T/G-rich light-strand gene (*ND6*). We analyzed all 13 genes under the `Vertebrate-mtDNA` genetic code across 27 mammalian genomes.
2. **Bacterial Replication Strand Asymmetry (*Borrelia* linear chromosome):** The linear chromosome of *Borrelia* replicates bidirectionally from a central origin, creating strong nucleotide skews across 13 Lyme and Relapsing Fever genomes. Leading-strand genes (*flaB*, *gyrA*, *gyrB*, *rpoB*) have positive GC skew and negative AT skew, whereas lagging-strand genes (*mutL*, *pyk*, *metK*, *p66*) have negative GC skew and positive AT skew.
3. **Mammalian Recombination Hotspots and gBGC:** Mammalian genes in high-recombination regions experience strong GC-biased gene conversion (*ADORA3*, *vWF*, and $\beta$-globin).
4. **Viral Host Deamination:** Retroviral and hepatitis genes edited by host cytidine (APOBEC3) or adenosine (ADAR) deaminases (HIV-1 *vif*, HIV-1 *RT*, and Hepatitis D virus).
5. **Empirical Genomic Controls and Deep Vertebrate Alignments (VGP & TOGA2):** Six reference alignments with known selection profiles (*PTS*, Encephalitis virus *env*, *rbcL*, *RBP3*, SARS-CoV-2 Spike, and *RH1*) plus 24 deep vertebrate coding alignments spanning 464 to 869 species (5,923 codons; Supplementary Table S4).

Across the 33 asymmetry alignments (14,194 codon sites; Supplementary Table S2), all 9 compositionally homogeneous ($K = 1$) alignments showed 100% concordance (0 discordant sites) between `FEL` and `dFEL`. Across the 24 heterogeneous ($K = 3$) asymmetry alignments, stationary `FEL` and directional `dFEL` differed at **81 codon sites** ($p \leq 0.05$): 39 in *Borrelia* leading/lagging strands, 28 in mammalian mtDNA, 11 in heterogeneous controls, and 3 in mammalian gBGC hotspots. Stationary `FEL` called 48 of these sites purifying that became neutral under `dFEL` once lineage-specific synonymous turnover was modeled, whereas `dFEL` unmasked 32 purifying sites where silent stasis had depressed $\hat{\alpha}_s$ under `FEL`. Similarly, across the 20 avian macro- and microchromosomal alignments (10,091 codons; Supplementary Table S1), all 10 homogeneous ($K = 1$) alignments showed 0 discordant sites, whereas the 10 heterogeneous ($K = 3$) alignments produced **95 discordant sites** ($p \leq 0.05$), tempering 67 `FEL`-only purifying calls and unmasking 4 additional diversifying calls on recombination-dense microchromosomes. Across all 176 discordant sites in the avian and asymmetry suites, `dFEL` reduced across-site $\mathrm{Var}(\hat{\alpha}_s)$ by 88.5%

**Table S1. Empirical benchmark comparison between Stationary FEL and Directional-FEL (dFEL).** Datasets encompass outgroup-free rooting benchmarks, viral strand-asymmetric genomes (Influenza A virus 8 segments), and avian macro- vs. microchromosomal genes evaluating the "Avian Paradox" (Bolívar et al. 2019). Site counts are evaluated at nominal significance $p \leq 0.05$.

| Benchmark System | Suite | Taxa | Codons | CP3 $\chi^2$ ($p$-value) | Auto-Root ($w$) | FEL (Pur / Div) | dFEL (Pur / Div) | Δ Calls |
|---|---|---|---|---|---|---|---|---|
| ***Rooting / Classical*** | | | | | | | | |
| Vertebrate beta-Globin | Vertebrata | 17 | 144 | 145.04 (1.1e-11) | `xenlaev` (4.7%) | 59 / 2 | 60 / 2 | **1** |
| Primate BRCA1 | Primates | 7 | 938 | 4.87 (0.999) | `Galago` (10.2%) | 38 / 0 | 38 / 0 | **0** |
| Vertebrate Rhodopsin RH1 | Vertebrata | 38 | 330 | 651.53 (2.1e-77) | `ZEBRAFINCH` (7.8%) | 222 / 1 | 228 / 1 | **6** |
| Bacterial PTS Trehalose Transporter | Bacteria | 16 | 639 | 221.14 (4.3e-25) | `NZ_AAFA0200000` (29.0%) | 287 / 0 | 286 / 0 | **1** |
| Drosophila Adh | Drosophila | 23 | 254 | 288.33 (1.5e-29) | `pic` (4.6%) | 123 / 1 | 123 / 1 | **0** |
| Mammalian RBP3 | Mammalia | 54 | 412 | 528.36 (3.4e-41) | `Pteropus` (4.4%) | 272 / 1 | 268 / 1 | **4** |
| ***Viral Strand Asymmetry*** | | | | | | | | |
| SARS-CoV-2 Spike | Coronaviridae | 180 | 1284 | 4.43 (1.000) | `BA_1_G339_S371` (0.3%) | 144 / 21 | 144 / 21 | **0** |
| Influenza A HA (H7) | Influenzavirus A | 146 | 572 | 540.01 (4.3e-04) | `Node99 - Node8` (0.5%) | 418 / 3 | 418 / 3 | **0** |
| Influenza A Matrix (MP) | Influenzavirus A | 327 | 253 | 771.37 (1.000) | `Node162 - Node` (0.2%) | 199 / 0 | 199 / 0 | **0** |
| Influenza A NA (N7) | Influenzavirus A | 92 | 472 | 239.7 (0.928) | `2005\|Avian\|A/A` (0.9%) | 323 / 2 | 323 / 2 | **0** |
| Influenza A Nucleoprotein (NP) | Influenzavirus A | 355 | 499 | 650.72 (1.000) | `2010\|Bat\|A/bat` (0.3%) | 430 / 2 | 430 / 2 | **0** |
| Influenza A Non-structural (NS) | Influenzavirus A | 340 | 233 | 301.7 (1.000) | `1963\|Avian\|A/t` (0.2%) | 139 / 17 | 139 / 17 | **0** |
| Influenza A Polymerase (PA) | Influenzavirus A | 332 | 717 | 938.06 (0.893) | `2009\|A/bat/Gua` (0.5%) | 608 / 0 | 608 / 0 | **0** |
| Influenza A Polymerase (PB1) | Influenzavirus A | 326 | 758 | 1244.45 (9.0e-09) | `2010\|Bat\|A/bat` (0.2%) | 655 / 1 | 655 / 1 | **0** |
| Influenza A Polymerase (PB2) | Influenzavirus A | 351 | 760 | 1198.11 (9.5e-04) | `1994\|Avian\|A/d` (0.2%) | 688 / 1 | 688 / 1 | **0** |
| ***Avian Macrochromosome*** | | | | | | | | |
| Avian ACBD7 (Macro) | Aves | 44 | 88 | 19.23 (1.000) | `Node41 - Node4` (1.2%) | 22 / 1 | 22 / 1 | **0** |
| Avian ACTR3 (Macro) | Aves | 44 | 330 | 15.64 (1.000) | `HLdicExi1` (1.2%) | 82 / 0 | 82 / 0 | **0** |
| Avian AASDH (Macro) | Aves | 44 | 1125 | 83.22 (0.999) | `HLcalPug1` (6.2%) | 291 / 12 | 291 / 12 | **0** |
| Avian ADAL (Macro) | Aves | 44 | 291 | 46.69 (1.000) | `HLamaAes1` (1.2%) | 106 / 2 | 106 / 2 | **0** |
| Avian ACPP (Macro) | Aves | 44 | 403 | 21.99 (1.000) | `HLdicExi1` (1.4%) | 105 / 5 | 105 / 5 | **0** |
| Avian ABCB5 (Macro) | Aves | 39 | 866 | 36.45 (1.000) | `HLlepAsp1` (2.1%) | 242 / 9 | 242 / 9 | **0** |
| Avian ABCE1 (Macro) | Aves | 44 | 599 | 18.74 (1.000) | `Node19 - Node1` (1.3%) | 214 / 1 | 214 / 1 | **0** |
| Avian ACTR3B (Macro) | Aves | 43 | 418 | 224.48 (1.6e-07) | `HLempTra1` (1.7%) | 154 / 1 | 155 / 1 | **1** |
| Avian ABHD13 (Macro) | Aves | 44 | 337 | 33.15 (1.000) | `HLlepAsp1` (1.5%) | 96 / 0 | 96 / 0 | **0** |
| Avian A1CF (Macro) | Aves | 44 | 574 | 1037.94 (7.6e-142) | `HLcinPul1` (42.4%) | 299 / 3 | 303 / 3 | **6** |
| ***Avian Microchromosome*** | | | | | | | | |
| Avian AANAT (Micro Chr28) | Aves | 41 | 205 | 198.35 (9.0e-06) | `HLserCan1` (5.0%) | 94 / 0 | 92 / 0 | **2** |
| Avian ABHD14A (Micro Chr26) | Aves | 39 | 242 | 79.85 (0.994) | `opiHoa1` (9.6%) | 69 / 0 | 69 / 0 | **0** |
| Avian ACKR1 (Micro Chr15) | Aves | 22 | 319 | 258.45 (1.7e-25) | `HLcalPug1` (20.5%) | 90 / 3 | 90 / 4 | **11** |
| Avian ACYP1 (Micro) | Aves | 40 | 99 | 71.26 (1.000) | `HLchaPel1` (2.3%) | 51 / 0 | 51 / 0 | **0** |
| Avian ACTC2L (Micro) | Aves | 34 | 375 | 335.78 (1.6e-27) | `HLserCan1` (13.6%) | 184 / 0 | 179 / 0 | **5** |
| Avian ACTG1 (Micro) | Aves | 34 | 375 | 335.78 (1.6e-27) | `HLserCan1` (14.2%) | 184 / 0 | 179 / 0 | **5** |
| Avian ACOT11 (Micro) | Aves | 43 | 561 | 214.38 (1.5e-06) | `opiHoa1` (6.1%) | 244 / 1 | 235 / 2 | **10** |
| Avian ACOT7 (Micro) | Aves | 44 | 324 | 157.87 (0.043) | `HLlepAsp1` (1.5%) | 161 / 0 | 163 / 0 | **2** |
| Avian AAR2 (Micro) | Aves | 26 | 336 | 118.78 (9.6e-04) | `HLserCan1` (10.2%) | 120 / 2 | 107 / 2 | **13** |
| Avian ABCA7 (Micro) | Aves | 38 | 2224 | 407.23 (2.0e-35) | `HLcolLiv2` (99.6%) | 877 / 9 | 855 / 11 | **40** |

(12.53 → 1.44) and reduced whole-gene Var($\hat{\alpha}_s$) by 62.6% in *RBP3*, 49.9% in *ACTG1* and *ACTC2L*, 42.9% in *ABCA7*, 41.8% in *AAR2*, and 30.9% in *RH1* (Supplementary Table S4, Part B). Finally, across the 24 deep vertebrate VGP and TOGA2 alignments (5,923 sites; Supplementary Table S4, Part A), `FEL` and `dFEL` agreed at 99.56% of sites ($p \leq 0.10$), with 100% agreement across all 14 homogeneous ($K = 1$) alignments and 26 discordant calls across the 10 heterogeneous ($K = 3$) alignments.

**Table S2. Comprehensive Known Asymmetry Benchmark: Stationary FEL vs. Directional dFEL Across Biological Arenas of Mechanistic Asymmetry.** Datasets represent biological systems with experimentally or genomic-mechanistically proven AT/GC composition skews: Vertebrate mtDNA heavy vs. light strand transcription, bacterial replication leading vs. lagging strand architecture (*Borrelia*), mammalian gBGC and recombination hotspots, viral host deamination, and empirical genomic controls. Discordant sites reflect shifts in site-level selective status at $p \leq 0.05$.

| | | | | Base Composition | | FEL ($p \leq 0.05$) | | dFEL ($p \leq 0.05$) | | |
|---|---|---|---|---|---|---|---|---|---|---|
| **Dataset** | **Suite** | **Taxa** | **Codons** | **CP3 $\chi^2$** | **GC3 Range (%)** | **Pur** | **Div** | **Pur** | **Div** | **Disc** |
| *Vertebrate mtDNA (H vs L)* | | | | | | | | | | |
| Mammalian mtDNA ND1 (Heavy Strand (A/C-rich)) | Vertebrate mtDNA ( | 27 | 319 | 344.7$^{***}$ | 22–52 | 273 | 0 | 273 | 0 | **2** |
| Mammalian mtDNA ND2 (Heavy Strand (A/C-rich)) | Vertebrate mtDNA ( | 27 | 350 | 348.9$^{***}$ | 22–52 | 262 | 0 | 259 | 0 | **3** |
| Mammalian mtDNA COX1 (Heavy Strand (A/C-rich)) | Vertebrate mtDNA ( | 27 | 523 | 585.0$^{***}$ | 23–50 | 493 | 0 | 493 | 0 | **2** |
| Mammalian mtDNA COX2 (Heavy Strand (A/C-rich)) | Vertebrate mtDNA ( | 27 | 231 | 304.5$^{***}$ | 23–54 | 207 | 0 | 207 | 0 | **2** |
| Mammalian mtDNA ATP8 (Heavy Strand (A/C-rich)) | Vertebrate mtDNA ( | 27 | 70 | 96.8 | 21–48 | 31 | 1 | 31 | 1 | **0** |
| Mammalian mtDNA ATP6 (Heavy Strand (A/C-rich)) | Vertebrate mtDNA ( | 27 | 226 | 281.9$^{***}$ | 15–48 | 195 | 0 | 196 | 0 | **1** |
| Mammalian mtDNA COX3 (Heavy Strand (A/C-rich)) | Vertebrate mtDNA ( | 27 | 262 | 264.8$^{***}$ | 23–50 | 232 | 0 | 233 | 0 | **1** |
| Mammalian mtDNA ND3 (Heavy Strand (A/C-rich)) | Vertebrate mtDNA ( | 27 | 117 | 183.4$^{***}$ | 19–47 | 96 | 0 | 98 | 0 | **2** |
| Mammalian mtDNA ND4L (Heavy Strand (A/C-rich)) | Vertebrate mtDNA ( | 27 | 99 | 133.5$^{***}$ | 24–57 | 79 | 0 | 80 | 0 | **1** |
| Mammalian mtDNA ND4 (Heavy Strand (A/C-rich)) | Vertebrate mtDNA ( | 27 | 459 | 514.4$^{***}$ | 21–51 | 384 | 0 | 380 | 0 | **6** |
| Mammalian mtDNA ND5 (Heavy Strand (A/C-rich)) | Vertebrate mtDNA ( | 27 | 616 | 562.0$^{***}$ | 22–54 | 469 | 1 | 469 | 0 | **5** |
| Mammalian mtDNA ND6 (Light Strand (Inverted T/G)) | Vertebrate mtDNA ( | 27 | 180 | 277.4$^{***}$ | 19–45 | 138 | 0 | 139 | 0 | **1** |
| Mammalian mtDNA CYTB (Heavy Strand (A/C-rich)) | Vertebrate mtDNA ( | 27 | 383 | 328.8$^{***}$ | 27–52 | 329 | 1 | 329 | 1 | **2** |
| *Bacterial Replication Asymmetry* | | | | | | | | | | |
| Borrelia flaB (Leading Strand) | Bacterial Replicat | 13 | 337 | 68.1$^{***}$ | 16–28 | 135 | 0 | 136 | 0 | **5** |
| Borrelia gyrA (Leading Strand) | Bacterial Replicat | 13 | 810 | 86.9$^{***}$ | 17–31 | 306 | 0 | 295 | 0 | **13** |
| Borrelia gyrB (Leading Strand) | Bacterial Replicat | 13 | 634 | 82.2$^{***}$ | 19–33 | 256 | 0 | 251 | 0 | **5** |
| Borrelia rpoB (Leading Strand) | Bacterial Replicat | 13 | 1442 | 346.0$^{***}$ | 16–31 | 651 | 0 | 649 | 0 | **6** |
| Borrelia mutL (Lagging Strand) | Bacterial Replicat | 13 | 624 | 117.2$^{***}$ | 17–30 | 165 | 0 | 167 | 0 | **6** |
| Borrelia pyk (Lagging Strand) | Bacterial Replicat | 13 | 477 | 36.2 | 17–26 | 211 | 0 | 211 | 0 | **0** |
| Borrelia metK (Lagging Strand) | Bacterial Replicat | 13 | 393 | 63.7$^{**}$ | 18–30 | 139 | 0 | 139 | 0 | **4** |
| Borrelia p66 (Lagging Strand) | Bacterial Replicat | 13 | 645 | 85.7$^{***}$ | 19–29 | 179 | 1 | 179 | 1 | **0** |
| *Mammalian gBGC & Isochores* | | | | | | | | | | |
| Mammalian ADORA3 (Isochore) | Mammalian gBGC | 67 | 107 | 140.2 | 54–84 | 56 | 1 | 56 | 1 | **0** |
| Mammalian vWF (Recomb Hotspot) | Mammalian gBGC | 62 | 392 | 923.0$^{***}$ | 69–95 | 250 | 2 | 248 | 2 | **2** |
| Mammalian Beta-Globin | Mammalian gBGC | 17 | 144 | 145.0$^{***}$ | 49–87 | 59 | 2 | 60 | 2 | **1** |
| *Viral Host Deamination* | | | | | | | | | | |
| HIV-1 Vif (APOBEC3 Target) | Viral Deamination | 29 | 192 | 8.0 | 33–38 | 22 | 8 | 22 | 8 | **0** |
| HIV-1 Reverse Transcriptase | Viral Deamination | 476 | 335 | 385.0 | 23–34 | 209 | 10 | 209 | 10 | **0** |
| Hepatitis Delta (Host Editing) | Viral Deamination | 33 | 196 | 28.6 | 66–73 | 49 | 8 | 49 | 8 | **0** |
| *Empirical Genomic Controls* | | | | | | | | | | |
| Bacterial PTS Transporter | Genomic Control | 16 | 639 | 221.1$^{***}$ | 25–38 | 287 | 0 | 286 | 0 | **1** |
| Encephalitis Virus Env | Genomic Control | 23 | 500 | 29.3 | 50–57 | 224 | 1 | 224 | 1 | **0** |
| Plastid rbcL | Genomic Control | 483 | 466 | 729.2 | 21–36 | 350 | 13 | 350 | 13 | **0** |
| Retinol Binding Protein 3 | Genomic Control | 54 | 412 | 528.4$^{***}$ | 63–90 | 272 | 1 | 268 | 1 | **4** |
| SARS-CoV-2 Spike | Genomic Control | 180 | 1284 | 4.4 | 26–27 | 144 | 21 | 144 | 21 | **0** |
| Vertebrate Rhodopsin RH1 | Genomic Control | 38 | 330 | 651.5$^{***}$ | 56–90 | 222 | 1 | 228 | 1 | **6** |

**Table S3. Supplementary Table S3: Head-to-head performance comparison between the Suzuki, Gojobori & Kumar (2009) counting method, Stationary FEL, and Directional dFEL.** Results compare the unweighted counting method (SG99; $1:1:1:1$), the CpG-adjusted counting method (SGK09; $40:4:4:1$), Stationary FEL, and Directional dFEL across Primate *PRM1* ($K = 1$), *Drosophila Adh* ($K = 3$), Avian *ABHD14A* ($K = 1$), and Avian *AAR2* ($K = 3$). Counts report diversifying / purifying sites identified at nominal significance levels $p \leq 0.05$ ($p \leq 0.10$).

| Dataset | Taxa | Codons | SG99 (1 : 1 : 1 : 1) Div / Pur ($p \leq 0.05/0.10$) | SGK09 (40 : 4 : 4 : 1) Div / Pur ($p \leq 0.05/0.10$) | Stationary FEL Div / Pur ($p \leq 0.05/0.10$) | **Directional dFEL Div / Pur** ($p \leq 0.05/0.10$) |
|---|---|---|---|---|---|---|
| Primate *PRM1* ($K = 1$) | 24 | 48 | 0 / 0 (1 / 0) | 0 / 0 (0 / 0) | 2 / 1 (3 / 2) | **2 / 1 (3 / 2)** |
| *Drosophila Adh* ($K = 3$) | 23 | 254 | 0 / 89 (0 / 107) | 0 / 69 (1 / 86) | 1 / 123 (2 / 142) | **1 / 123 (1 / 145)** |
| Avian *ABHD14A* ($K = 1$) | 39 | 242 | 0 / 62 (5 / 68) | 3 / 44 (6 / 62) | 0 / 69 (2 / 86) | **0 / 69 (2 / 86)** |
| Avian *AAR2* ($K = 3$) | 26 | 336 | 0 / 63 (3 / 76) | 10 / 55 (15 / 76) | 2 / 120 (4 / 142) | **2 / 107 (6 / 135)** |

**Table S4. Supplementary Table S4: Deep Vertebrate Phylogenomics Benchmark (VGP and TOGA2 Cohorts, 464–869 Taxa) and Whole-Gene Synonymous Rate Variance (Var($\hat{\alpha}_s$)) Reduction.** Part A compares stationary `FEL` and directional `dFEL` ($p \leq 0.10$) alongside `MEME` ($p \leq 0.05$) across 12 Vertebrate Genomes Project (VGP) and 12 TOGA2 coding alignments (5,923 codons total). In all 14 genes with $K = 1$ stationary regime, `FEL` and `dFEL` show 100% concordance (0 discordant sites); all 26 discordant sites occur in the 10 compositionally heterogeneous ($K = 3$) loci. Part B summarizes whole-gene across-site synonymous rate dispersion (SD($\hat{\alpha}_s$) and $\Delta$Var($\hat{\alpha}_s$)) in representative compositionally skewed loci.

| Cohort | Gene | Taxa | Codons | Regimes ($K$) | `FEL` Pur | `FEL` Div | `dFEL` Pur | `dFEL` Div | `MEME` Div | Discordant | Auto-Root Check | Key Site Concordance & Directional Shifts |
|---|---|---|---|---|---|---|---|---|---|---|---|---|
| ***Part A: Deep Vertebrate Alignments (VGP & TOGA2, 24 Loci, 5,923 Codons)*** | | | | | | | | | | | | |
| VGP | *VAMP3* | 538 | 105 | 3 | 77 | 3 | 77 | 1 | 3 | 2 | 1,001 br (0 diff) | Removes 2 false Div: site 91 (3 2-hit bursts) & site 94 (15 I → V [GC ↑], `MEME` $p = 0.372$). |
| VGP | *RPS20* | 830 | 138 | 3 | 114 | 0 | 114 | 0 | 1 | 0 | 1,521 br (0 diff) | 100% site concordance across 830 vertebrate taxa. |
| VGP | *PSMA4* | 843 | 264 | 3 | 249 | 0 | 249 | 0 | 0 | 0 | — | 100% site concordance under strong purifying constraint. |
| VGP | *MS4A12* | 464 | 286 | 1 | 80 | 35 | 80 | 35 | 52 | 0 | — | Homogeneous ($K = 1$); 100% site concordance (35 Div, 80 Pur). |
| VGP | *ADORA3* | 740 | 352 | 1 | 232 | 9 | 232 | 9 | 10 | 0 | 1,442 br (0 diff) | Homogeneous ($K = 1$); ML root placed on `eleEdw1` (Afrotheria). |
| VGP | *ZPBP* | 804 | 423 | 3 | 245 | 19 | 244 | 20 | 23 | 2 | — | Detects Div site 95 ($9 \times$ S → A [GC ↑]); removes false Pur site 97 ($9 \times$ G → E [AT ↑]). |
| VGP | *KRT20* | 562 | 497 | 3 | 247 | 24 | 250 | 24 | 42 | 3 | — | Detects 3 Pur sites (343, 385, 437) masked by bidirectional GC/AT flux. |
| VGP | *RTN4R* | 723 | 499 | 3 | 433 | 0 | 434 | 1 | 9 | 4 | — | Detects Div site 319 ($p = 0.052$) & 2 Pur (291, 313); removes 2-hit false Pur site 370. |
| VGP | *CD4* | 724 | 517 | 1 | 203 | 42 | 203 | 42 | 75 | 0 | — | Homogeneous ($K = 1$); 100% site concordance (42 Div, 203 Pur). |
| VGP | *MYOC* | 688 | 546 | 3 | 368 | 10 | 369 | 12 | 23 | 5 | — | Detects Div sites 15 (`MEME` $p = 0.048$) & 98, 2 Pur (32, 90); removes 2-hit false Pur site 218 (`MEME` $p = 0.004$). |
| VGP | *TNFRSF1A* | 548 | 584 | 3 | 255 | 23 | 253 | 25 | 40 | 6 | — | Detects Div sites 54 & 367, Pur site 538; removes 3 false Pur (113 [`MEME` $p = 0.009$], 360, 382). |
| VGP | *DMRT3* | 575 | 601 | 3 | 424 | 0 | 427 | 0 | 1 | 3 | — | Detects 3 Pur sites (492, 538, 584) after normalizing directional GC/AT transitions. |
| TOGA2 | *TIMM10* | 859 | 102 | 1 | 83 | 0 | 83 | 0 | 1 | 0 | 1,262 br (0 diff) | Homogeneous ($K = 1$); 100% site concordance between `keep` and `auto` root. |
| TOGA2 | *WDR83OS* | 835 | 104 | 1 | 93 | 0 | 93 | 0 | 1 | 0 | 1,232 br (0 diff) | Homogeneous ($K = 1$); ML root independently placed on `eleEdw1` (Afrotheria). |
| TOGA2 | *NPY* | 860 | 114 | 1 | 78 | 3 | 78 | 3 | 6 | 0 | 1,290 br (0 diff) | Homogeneous ($K = 1$); 100% site concordance (3 Div, 78 Pur). |
| TOGA2 | *LSM7* | 848 | 138 | 3 | 99 | 0 | 100 | 0 | 2 | 1 | — | Resolves boundary stall at site 124 (`FEL` $p = 1.00$ → `dFEL` Pur $p = 0.007$). |
| TOGA2 | 8 Other Loci† | 839–869 | 1,025 | 1 | 801 | 3 | 801 | 3 | 24 | 0 | — | All 8 homogeneous ($K = 1$) alignments show 100% site concordance (0 discordant). |
| **Total (24 Loci)** | — | **464–869** | **5,923** | **1–3** | **3,881** | **171** | **3,887** | **175** | **313** | **26** | **6/6 (100% inv.)** | **99.56% overall concordance; 100% on $K = 1$ loci; 26 shifts on $K = 3$ loci.** |
| ***Part B: Coupling of Directional Drift and Apparent Synonymous Rate Variation (Var($\hat{\alpha}_s$))*** | | | | | | | | | | | | |
| Avian + Asym. | Discordant Sites | 10–180 | 176 | 3 | 116 | 3 | 51 | 6 | — | 176 | — | SD($\hat{\alpha}_s$) : 3.540 → 1.199 (Var($\hat{\alpha}_s$) : 12.53 → 1.44, −**88.5**% across-site SRV reduction). |
| Mammalian | *RBP3* (Whole Gene) | 32 | 412 | 3 | 272 | 1 | 268 | 1 | — | 4 | — | SD($\hat{\alpha}_s$) : 1.852 → 1.132 (CV : 1.52 → 0.99, −**62.6**% whole-gene Var($\hat{\alpha}_s$) reduction). |
| Avian Micro | *ACTG1* & *ACTC2L* | 37 | 750 | 3 | 368 | 0 | 358 | 0 | — | 10 | — | SD($\hat{\alpha}_s$) : 5.948 → 4.209 (CV : 2.21 → 1.86, −**49.9**% whole-gene Var($\hat{\alpha}_s$) reduction). |
| Avian Micro | *ABCA7* (Whole Gene) | 40 | 2,224 | 3 | 877 | 9 | 855 | 11 | — | 40 | — | SD($\hat{\alpha}_s$) : 4.078 → 3.081 (CV : 2.38 → 2.08, −**42.9**% whole-gene Var($\hat{\alpha}_s$) reduction). |
| Avian Micro | *AAR2* (Whole Gene) | 26 | 336 | 3 | 120 | 2 | 107 | 2 | — | 13 | — | SD($\hat{\alpha}_s$) : 3.527 → 2.689 (CV : 1.75 → 1.79, −**41.8**% whole-gene Var($\hat{\alpha}_s$) reduction). |
| Vertebrate | *RH1* (Whole Gene) | 38 | 330 | 3 | 222 | 1 | 228 | 1 | — | 6 | — | SD($\hat{\alpha}_s$) : 2.580 → 2.145 (CV : 1.86 → 1.51, −**30.9**% whole-gene Var($\hat{\alpha}_s$) reduction). |
| *Drosophila* | *Adh* (Whole Gene) | 23 | 254 | 3 | 123 | 1 | 123 | 1 | — | 0 | — | SD($\hat{\alpha}_s$) : 8.111 → 7.567 (CV : 3.62 → 3.39, −**13.0**% whole-gene Var($\hat{\alpha}_s$) reduction). |
| Avian Micro | *AANAT* (Whole Gene) | 33 | 205 | 3 | 94 | 0 | 92 | 0 | — | 2 | — | SD($\hat{\alpha}_s$) : 5.479 → 5.126 (CV : 2.41 → 2.64, −**12.5**% whole-gene Var($\hat{\alpha}_s$) reduction). |

†The 8 additional homogeneous ($K = 1$) TOGA2 loci are *DYNLL1* (120 codons), *NDUFA2* (124), *CNFN* (126), *ATP6V1F* (129), *C12orf57* (131), *ANAPC15* (133), *RPL27* (138), and *COX6A1* (144).
